# Context-Dependent Design of Induced-fit Enzymes using Deep Learning Generates Well Expressed, Thermally Stable and Active Enzymes

**DOI:** 10.1101/2023.07.27.550799

**Authors:** Lior Zimmerman, Noga Alon, Itay Levin, Anna Koganitsky, Nufar Shpigel, Chen Brestel, Gideon D. Lapidoth

## Abstract

The potential of engineered enzymes in practical applications is often constrained by limitations in their expression levels, thermal stability, and the diversity and magnitude of catalytic activities. *De-novo* enzyme design, though exciting, is challenged by the complex nature of enzymatic catalysis. An alternative promising approach involves expanding the capabilities of existing natural enzymes to enable functionality across new substrates and operational parameters. To this end we introduce CoSaNN (Conformation Sampling using Neural Network), a novel strategy for enzyme design that utilizes advances in deep learning for structure prediction and sequence optimization. By controlling enzyme conformations, we can expand the chemical space beyond the reach of simple mutagenesis. CoSaNN uses a context-dependent approach that accurately generates novel enzyme designs by considering non-linear relationships in both sequence and structure space. Additionally, we have further developed SolvIT, a graph neural network trained to predict protein solubility in *E.Coli*, as an additional optimization layer for producing highly expressed enzymes. Through this approach, we have engineered novel enzymes exhibiting superior expression levels, with 54% of our designs expressed in E.Coli, and increased thermal stability with more than 30% of our designs having a higher Tm than the template enzyme. Furthermore, our research underscores the transformative potential of AI in protein design, adeptly capturing high order interactions and preserving allosteric mechanisms in extensively modified enzymes. These advancements pave the way for the creation of diverse, functional, and robust enzymes, thereby opening new avenues for targeted biotechnological applications.

## Introduction

Natural evolution in proteins introduces variability by employing mechanisms such as mutations, gene duplication, and homologous recombination, which often result in the emergence of novel structures and functions ^1–3^. This phenomenon is most evident in multidomain proteins, where independently folding domains are fused together to generate new protein structure and function. Not surprisingly, these principles are not restricted to multi-domain proteins but also function within individual domains, giving rise to novel globular protein folds by blending secondary and supersecondary elements^4–6^.

This phenomenon has been exploited in the past to generate new protein variants through techniques such as DNA shuffling ^7^ to create novel β-lactamase enzymes. Later work expanded on this technique to combine random *E.coli* DNA fragments to generate novel protein folds ^8^, demonstrating that structural elements can be imported from distant and even unrelated protein folds. The concept of fragment recombination has been further refined to include structural information, thereby minimizing structural disruption and increasing the likelihood of generating functional proteins ^9^. Similar approaches have been successfully used to combine sub-domain fragments from different protein folds, demonstrating that even distantly related proteins can be recombined to generate novel, functional proteins ^10^.

These methods, dependent on rational segmentation or random fragmentation and fusion of parent enzymes, have limited predictive capacity for the resulting output. Instead, they hinge on extensive high-throughput screening, requiring multiple iterative cycles of experimentation.

Building upon these engineering strategies, computational protein design contributes to the field by not only selecting fragments for recombination but also modeling the resulting novel conformations and optimizing their amino-acid sequences in-silico. Prominent examples for this approach are the SEWING algorithm developed by Jacobs *et al*.^11^, and AbDesign developed by Lapidoth *et al.* . Both algorithms were developed within the RosettaDesign software suite^14^ environment and use combinatorial backbone assembly sampled from natural protein conformation library.

However, these methods have some shortcomings. Primarily, the way the “donor” protein fragment structurally aligns within the protein scaffold is dependent on how well the ’anchor regions’ of the fragment and scaffold match up. This process is vulnerable to what is known as the ’lever effect’: slight variations in the anchor regions, stemming from the fragment and scaffold protein alignment, can cause substantial shifts in regions distant from the anchor, leading to inaccurate backbone conformations. Additionally, it is well established that protein fragments can adopt alternative conformation when placed in new environmental contexts^15,10,16^. Yet, both algorithms mentioned earlier make the assumption that the sampled fragments remain rigid and maintain their original conformation within the context of the new scaffold. Lastly, both methods use RosettaDesign to design the sequence of the entire protein to optimize its conformation. RosettaDesign uses a Monte Carlo-based algorithm to search the sequence space for low-energy sequences for a given protein backbone. Other than being very computationally intensive, its energy function does not capture all aspects of protein thermodynamics and kinetics accurately ^17^. The requirement for Rosetta’s energy function to be pairwise decomposable means that higher order interactions between residues are missed. This can generate protein designs that, while having favorable Rosetta energies (analogous to folding energy) do not fold well in reality. Moreover, this presents a particular challenge when attempting to computationally design allosteric proteins^18^. Allosteric proteins require the stabilization of multiple distinct, functionally relevant conformations, a complexity not easily captured by pairwise energy computations.

Here we present CoSaNN (Conformation Sampling using Neural Network), a design strategy for creating novel enzyme conformations that takes into account the surrounding structural and chemical context in which the fragment is located. The algorithm leverages the latest advances in Deep-learning based structure prediction, specifically DeepMind’s AlphFold2^19^. CoSaNN begins with a template structure. The structure is then segmented along structurally conserved points within the protein family. Unlike previous backbone sampling algorithms that require computationally intensive sampling of protein structures ^12^, new conformations are generated by creating sequence based chimeras, swapping sequence segments in the scaffold enzyme with sequence segments from donor proteins. The chimeric sequences are then modeled using AlphaFold.

AlphaFold and other similar neural network (NN) structure prediction methods are trained to map sequences to their corresponding structures. Through this process, these models effectively learn to recognize sequence-structure pattern motifs. In practice, this means that if a sequence fragment from protein A is inserted into an equivalent position in a homologous protein B, AlphaFold is capable of accurately folding the segment such that the conformation of both accepting template and grafted segment changes only slightly. We hypothesize that this ability is attributed to the contextual dependency learned by the model during training. This feature explains why when presented with sequences containing mutations known to cause experimental misfolding, AlphaFold tends to predict the ’native’ or typical fold.

In our approach, we utilize the context-dependent folding propensity of AlphaFold as an asset to generate realistic, native-like backbone conformations for our sequence chimeras. Upon generation of these conformations, we further optimize them through sequence design. This process aims to enhance the likelihood of the generated backbones folding correctly. For the sequence optimization step we chose to use the newly developed ProteinMPNN ^20^ as well as RosettaDesign ^14^. This choice of ProteinMPNN over the more conventional RosettaDesign aligns with our overarching design approach, which advocates for the utilization of context-dependent capabilities of neural networks.

ProteinMPNN is trained on a large dataset of protein structures to predict sequences that are compatible with a given protein backbone. This approach allows ProteinMPNN to capture higher order, non-linear relationships between sequence and structure than are explicitly included in the Rosetta energy function.

In our sequence design step, we chose to keep the catalytic residues fixed. This decision was informed by previous design iterations, which revealed a propensity for both RosettaDesign and ProteinMPNN to occasionally mutate these residues. This tendency is not unexpected, given the well-documented tradeoff between enzymatic functions and protein stability ^21^. It is worth noting that both algorithms are primarily optimized for ensuring foldability, not preserving catalytic function.

Moving forward, the further refinement and development of ProteinMPNN-like algorithms could potentially alleviate this dilemma by integrating catalytic function preservation into their optimization objectives, alongside foldability.

Lastly, the generated sequences are modeled again using AlphaFold and are ranked using a novel graph neural network classifier we developed, trained to predict soluble expression in *E.coli*. We developed this classifier specifically to address the challenges associated with predicting expression titers. This provides another layer of optimization, which is not explicitly accounted for in either ProteinMPNN or RosettaDesign.

In the present study, our method has been applied towards the design of enzymes that exhibit significant conformational changes upon substrate binding. This scenario poses a rigorous test for the robustness of the design strategy we have developed. Not only do we obtain highly active designs, without requiring laboratory evolution, but we maintain the fold’s allosteric mechanism, clearly demonstrating that our design approach accounts for long range, high order interactions both in sequence and structure space. We also demonstrate that our approach works well not only in scenarios that involve the grafting of a single segment, but also in grafting of multiple different segments, some are radically different than the acceptor segment; Finally, we show that our classifier based ranking system significantly increases the likelihood of finding well expressed proteins from the millions of generated in-silico designs.

## Methods

### Computational

#### Selection of template

Polyphosphate glucokinase (ppgmk) from Arthrobacter sp. (strain KM, Uniprot Id: Q7WT42) was selected as the template on which all backbone segments are grafted onto. The 3D structure of the protein was previously determined by X-Ray crystallography^22^ in the bound conformation with glucose and a pair of phosphates adjacent to the glucose substrate. (PDB Id: 1WOQ)

#### Selection of anchor positions

To determine the most structurally invariant anchors across the entire protein fold, we developed an algorithm that creates distance maps for all proteins within the same PFam fold as ppgmk (PF00480), clustered to 50% sequence identity. The distance maps are then modified to only include structurally conserved residues across the fold family, generating a ‘consensus’ distance map of the fold family. To compute the distance map, we employed mTm-Align^23^ to create a multiple structural alignment, reminiscent of the process for multiple sequence alignment. From the produced alignment, we derived a consensus that consisted only of positions uniformly aligned across all candidates. This consensus alignment informed the assembly of the consensus distance map, incorporating solely the residues present within the consensus. To identify invariant spatial position pairs across all family members, we measured the variance across all matching entries in the consensus distance map across all proteins. Positions showcasing the least variance were recognized as invariant and potentially suitable for accepting foreign segments.

The following anchor positions were used in this work: residues 81,122 for the first fragment and residues 159,194 for the second fragment. These positions were selected based on their spatial invariance across the entire fold family and based on their contribution to substrate recognition and catalysis ^24^.

#### Generating chimeric enzymes from segments of homologous proteins

Anchor positions were derived by obtaining the consensus anchor points (as described above) and mapping them to the template and donor proteins. The amino-acid sequence between the two anchors was then extracted for each of the entries in the Pfam fold family of ppgmk (PF00480) and inserted into the template sequence, replacing the original template’s equivalent sequence.

#### Modeling chimeric enzymes

Chimeric enzymes were modeled using the ColabFold^25^ implementation of AlphaFold2. The multiple sequence alignments required by AlphaFold2 as additional input were generated using mmseqs ^26^.

#### Sequence design of modeled enzymes

Sequence optimization for the chimeric enzymes was performed using either ProteinMPNN or RosettaDesign. A Position-Specific Scoring Matrix (PSSM) was utilized to effectively constrain the sequence space in both cases. For ProteinMPNN, the PSSM was incorporated as a bias to guide the model’s logit predictions. Conversely, for RosettaDesign, we used the PSSM-generated log-likelihood values to eliminate any amino acids that yielded log-likelihood ratio values less than zero.

#### Training an E.coli based heterologous expression predictor

The ESol dataset^27^ consisting of 3,173 translated E.coli proteins with their respective soluble fraction titers was downloaded from the author’s website. The structure of each protein in the dataset was modeled using AlphaFold2. Subsequently, a graph representation of the protein was created from the structure such that the protein’s amino acids form the graph vertices and an edge exists between two amino acids if the distance between them is < 5Å. 20% of the samples were set a side to serve as the test set while the rest of the samples were partitioned to 5 identically sized sets for a 5-fold cross validation training process. The solubility classifier is an ensemble of 5 models trained on the 5 folds. Each of the models is composed of two graph attention layers ^28^ and a ReLU activation function.

Following the attention layers, the graph representation is pooled by a Graph Multiset Transformer pooling operator ^29^. Finally, a linear layer is applied, followed by a sigmoid activation function. The implementation of the network was done in PyTorch^30^ and PyTorch-Geometric^31^.

#### Protein expression

NiCo21(DE3) E.coli competent cells (New England Biolabs, Ipswich, MA) were transformed using the heat shock method with a pET-28a plasmid that was cloned with our designed enzymes, with an addition of an N-terminal 6xHis-tag. The transformation was conducted in a 96-well plate (Cat: 701011, Wuxi-Nest, Jiangsu, China). After transformation, the cells were cultivated in their respective wells at 37℃ for 16 hours, in a 2xYT medium supplemented with 30µg/ml of kanamycin for selection. The cells were subsequently diluted in a 1:60 ratio into a deep-well 96-plate (Labcon, USA, Cat: LC 3909-525), reaching a final volume of 0.5ml per well. This dilution was made using fresh 2xYT medium containing 30µg/ml of kanamycin. The cultures were then incubated at 37℃ until they attained an absorbance of 0.6 at 600nm, indicating optimal growth.

Protein expression was initiated by the addition of isopropylthio-β-galactoside (IPTG) at a concentration of 0.2mM. The cultures were then induced for approximately 20 hours at 20℃. Following this induction period, the cells were harvested through centrifugation at 4,000g at a temperature of 4°C for 15 minutes

#### Protein isolation and quantification

The cells were resuspended in 100µl Qproteome Bacterial Protein Prep kit (Qiagen, Germany, cat. 37900). 1mg/ml lysozyme, 1:100 (v/v) Protease Inhibitor Cocktail (EDTA-Free, 100X in DMSO) (APExBIO, USA, Cat. K1010) and 1:1000 (v/v) benzonase were add to the lysis buffer. After 30 minutes on ice, the lysate was centrifuged (4,500 g; 4°C; 30 min) and the soluble fraction was incubated in a shaker with Ni-NTA Charged MagBeads (GeneScript, USA, cat. L00295) for 1h at 25℃. Prior to incubation with cells supernatant, the beads were pre-washed with a binding buffer (50mM TRIS-HCl pH 8.0, 300mM KCl, 0.1Mm ZnCl_2_ and 10mM imidazole). Subsequently, the beads were washed three times with 200µl wash buffer (50mM TRIS-HCl pH 8.0, 300mM KCl, 0.1Mm ZnCl_2_ and 30mM imidazole). His-tagged proteins were eluted with 50µl elution buffer (50mM TRIS-HCl pH 8.0, 100mM KCl, 0.1Mm ZnCl_2_ and 500mM imidazole). The eluted proteins concentrations were then inspected using standard SDS PAGE with Coomassie staining, and quantified using Coomassie Plus Bradford reagent Assay kit (Thermo Scientific, USA, cat. 23236).

#### Protein thermal stability

Protein thermal stability was assayed using Applied Biosystems, Protein Thermal Shift Dye Kit according to the manufacturer’s protocol and analyzed using Bio-Rad C1000 Touch Thermal cycler machine and software. The Sypro-Orange dye was excited at 470 nm and change to the fluorescence emission at 570 nm (Relative Fluorescence Unit-RFU) was measured in temperatures ranging from 25℃ to 95℃ with an increase of 1℃ per minute.

#### End-point Glucokinase activity

Endpoint enzymatic activity was assayed at 37°C unless indicated otherwise. The reaction mixture contained 50 mM Tris-HCl buffer (pH 8.0), 100 mM NaCl, 0.1 mM ZnCl2, 1.6 µg/ml polyphosphate, and 100 mM glucose. The reactions were initiated by the addition of the enzyme and terminated by freezing after 6 minutes. To determine the concentration of glucose-6-phosphate, PicoProbe™ Fructose-6-Phosphate Assay Kit (GenWay Biotech, Inc.), was used as instructed by the manufacturer with minor adaptations: 1) glucose-6-phosphate was used as a standard and 2) .

Glucose-6-phosphate isomerase was omitted from the reaction. Fluorescence was measured in a synergy HT-T1 plate reader (Biotek, USA), Ex. 528, Em. 590. The optimal temperature was determined by running an activity assay at varying temperatures between 25°C-95°C for 6 minutes.

In order to evaluate the enzymes’ stability at different temperatures, the enzymes were incubated at varying temperatures ranging from 25°C-80°C for indicated times (0-22 hours) in the enzymatic reaction buffer. Then, the temperature was changed to 37°C, glucose and polyphosphate were added, and the enzymatic activity was measured.

#### Glucokinase Kinetic Measurements

Enzyme kinetics were assayed for 20 minutes at 37°C using flat bottom UV-transparent plates (UV-Star 96 wells, Greiner, Bio-One #655801). The reaction mixture contained 50 mM Tris-HCl buffer (pH 8.0), 100 mM NaCl, 0.1 mM ZnCl2, 10 mM MgCl2, 66 µM NADP, and Converter Mix containing glucose 6-phosphate dehydrogenase from PicoProbe™ Fructose-6-Phosphate Assay Kit that was diluted 1500 fold . Km and Vmax values for glucose were determined by varying the glucose concentration from 0 to 166 mM while keeping the concentration of the polyphosphate at 8ug/ml. Km and Vmax values for polyphosphate were assayed by varying the polyphosphate concentration from 0 to 320µg/ml while keeping the concentration of glucose at 33mM. The formation of glucose-6-phosphate was monitored by coupling with 1500 fold dilution of glucose-6-phosphate dehydrogenase Converter Mix as mentioned above and measuring the formation of NADH at 340nm. Kinetic parameters were calculated by measuring the initial rates of reactions and fitting the data to Michaelis–Menten model using GraphPad Prism version 9.4.0 for Windows, (GraphPad Software, USA).

## Results

### The ROK carbohydrate kinase

The ROK fold family is an ideal testbed for our design method’s generality due to its complex conformational dynamics^32^ and functional diversity^33^. The ROK (Repressor, Open reading frame, Kinase) protein family (Pfam 00480) represents a functionally diverse collection of polypeptides that includes carbohydrate-dependent transcriptional repressors, sugar kinases, and yet uncharacterized open reading frames^34^. The ROK family is a member of the Actin-ATPase clan (Pfam CL 0108) that includes 21 distinct members, many of which couple phosphoryl transfer/hydrolysis to a functionally significant conformational change. The ROK fold consists of an N-terminal small α/β domain and a C-terminal large α/β domain, separated by a deep cleft that contains the active site, with the carbohydrate held in position by amino acid side chains from both domains^35^ (Fig. 1A). Binding of the carbohydrate causes a substantial rearrangement of the two domains, and the ATP binding site forms only after the carbohydrate binds, forming a closed catalytically active conformation (Fig. 1B). Kinetic analysis indicates that the enzymes show an ordered Bi–Bi sequential mechanism where the carbohydrate binds first, followed by ATP^32^.

**Figure 1.**
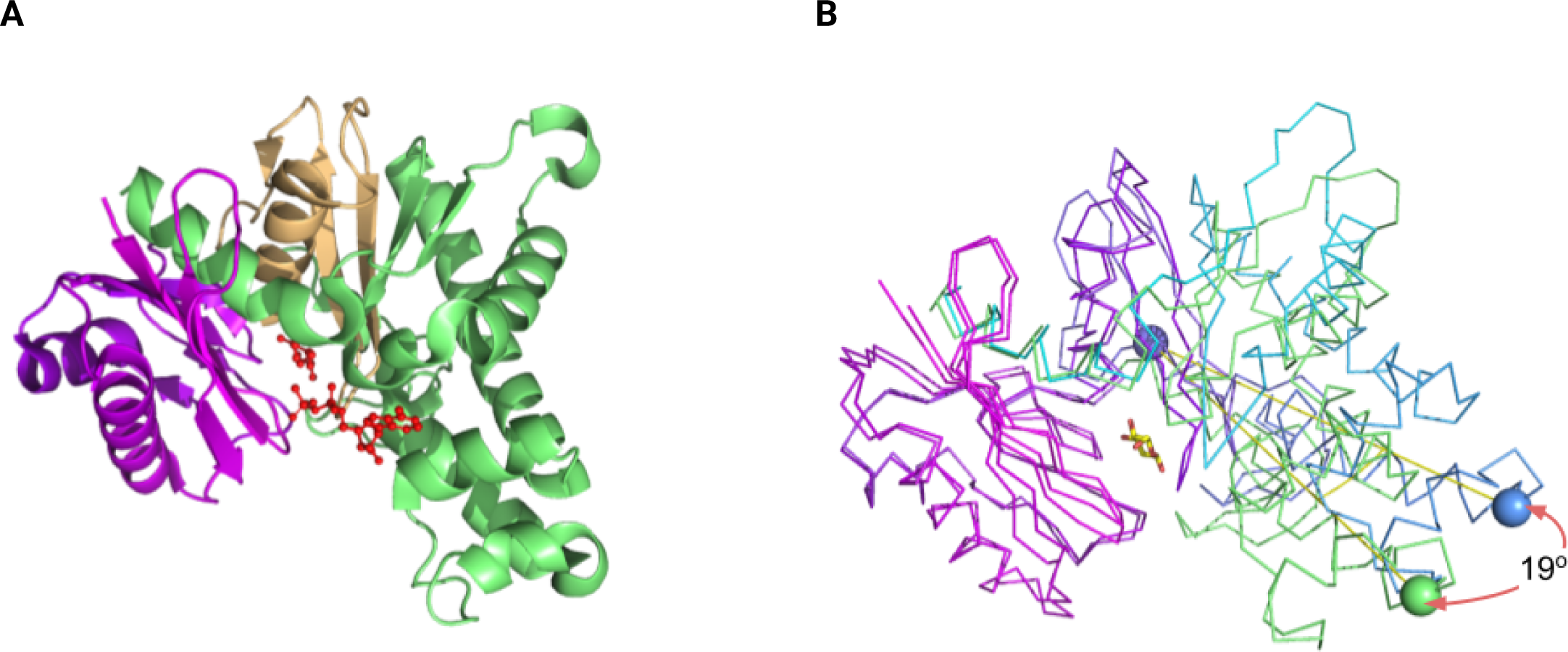
Structural features and mechanism of ROK kinases. (A) The ROK fold comprises an N-terminal small α/β domain (magenta), a C-terminal large α/β domain (green), and a hinge region (yellow), which together form a deep active-site cleft where the carbohydrate and ATP ligands (depicted as red ball-and-stick models) are positioned. (B) A significant conformational change occurs upon substrate binding, causing a 19-degree shift in the interdomain angle.

Few ROK kinases have been extensively characterized for their substrate specificity. Those that have been studied are selective for a narrow range of related sugars, with little activity against other sugars ^33^. Consistent with this, phylogenetic analysis indicates that sugar specificity is mainly encoded by sequence motifs found on the small N-terminal α/β domain ^24^.

While most ROK kinases characteristically prefer ATP as a phosphate donor, and require a magnesium ion ^24^, some of these enzymes can additionally utilize polyphosphate as a phosphate donor ^36^. Notably, the unique inorganic polyphosphate/ATP-glucomannokinase from Arthrobacter sp. strain KM (ppgmk) is the only inorganic phosphate independent ROK kinase whose structure has been determined (PDB 1WOQ)^22^. Ppgmk lacks two secondary structure motifs that are characteristic in the ROK family: the “ROK motif” (CXCGXXGC) which acts as a zinc finger motif, and a helical motif that clamps the adenosine substrate. The zinc finger motif functions as a substrate recognition mechanism, directing a conserved Histidine residue to form a hydrogen bond with the sugar substrate. This interaction triggers an allosteric conformational transformation that enables ATP binding in a bi-bi mechanism. Accordingly, ROK family members lacking this motif do not display the same bi-bi mechanism^32, 37^.

Another distinct feature of ppgmk is the phosphate binding mode. The crystal structure was determined with two phosphate ions bound at the boundary between the two domains, suggesting that polyphosphate binds between the two domains placing the terminal phosphate at a position amenable for phosphorylation (Fig. 2B).

**Figure 2.**
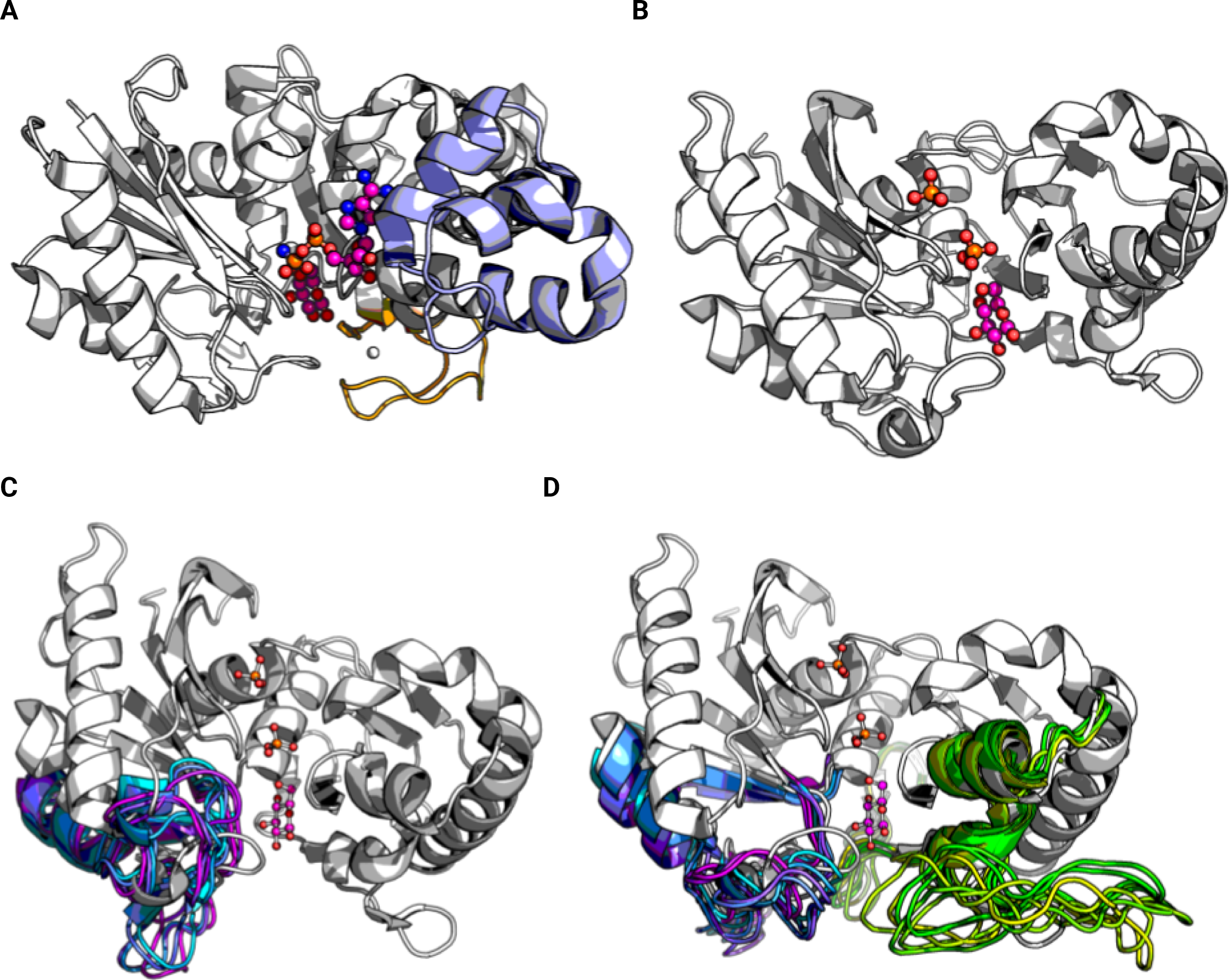
Comparison between ATP dependent and polyphosphate dependent ROK family sugar kinase and depiction of backbone sampling. (**A**) The structures of the ATP-dependent glucokinase from from Streptomyces griseus complexed with glucose (orange spheres) and ATP (pink spheres) (PDB ID: 3VGL) and (**B**) the inorganic Polyphosphate/ATP-Glucomannokinase (ppgmk) From Arthrobacter sp. complexed with glucose (yellow spheres) and phosphate ions (orange and red spheres). Ppgmk lacks the zinc-finger motif (colored green and yellow) and an additional secondary structure (colored tiel) that ‘clamps’ the ATP adenosine moiety. (**C)** Representative conformations of the N-terminal segment (colored blue-magenta) grafted onto ppgmk (colored white) created using Design Strategies 1&2. (**D**) Representative conformations of the N-terminal segment (colored blue-magenta) and the C-terminal segment (colored green-yellow) grafted onto ppgmk (colored white) created using Design Strategies 3 and 4. The glucose substrate and phosphate ions are shown as magenta and red spheres and orange and red spheres respectively.

Given the structural distinctiveness of the ROK ppgmk, suggesting also a unique activity mechanism, we decided to use it as the template onto which other segments would be grafted to, offering a challenging test case for our design methodology.

### Comparing design strategies

To ensure a robust and comprehensive evaluation of our design strategy, CoSaNN, we separated the design protocol into four distinct elements. These include two design tasks—backbone conformation sampling and sequence optimization—and two design strategies used for these tasks—neural network (NN)-based methods and Rosetta-based methods. Additionally, we tested both single segment replacement as well as multiple segment replacements (See Table 1). This comparative approach provides a meaningful assessment of each method’s strengths and weaknesses, allowing us to evaluate the overall effectiveness of our design strategy in a variety of conditions. Furthermore, by segregating the design tasks, we can more accurately pinpoint areas of success and necessary improvement within each procedural element.

**Table 1:** Computational Design Strategies.

| Design Strategy | Grafted segment | Backbone grafting method | Sequence design method | # of designs generated |
| --- | --- | --- | --- | --- |
| 1 | 81-122 | Rosetta based <sup>13</sup> | Rosetta Design | 40 |
| 2 | 81-122 | AlphaFold Sequence grafting | Rosetta Design | 119 |
| 3 | 81-122, 159-194 | AlphaFold Sequence grafting | Rosetta Design | 95 |
| 4 | 81-122, 159-194 | AlphaFold Sequence grafting | ProteinMPNN | 94 |

We decided to focus our design on two regions within the protein scaffold: a segment stretching residues 81-122 and the second stretching between residues 159-194 (numbering corresponding to the template protein ppgmk). The 81-122 segment is part of the N-terminal small domain and is largely where the sugar specificity motif lies^24^ (Fig. 2C). Particularly within ppgmk’s context, it contains a higher density of positively charged residues. These residues play a pivotal role in stabilizing the negatively charged polyphosphate molecule, a characteristic that appears to be less common among other ATP-dependent members of the ROK family. The 159-194 segment contains the characteristic zinc finger motif shared by nearly all members of the ROK family (Fig. 2D).

Intriguingly, this motif is absent from our template enzyme, further underscoring the unique structural characteristics that differentiate it from other members of the ROK family.

In the following section we elaborate on the characteristics of each design strategy.

### Design Strategy 1

As a control set we opted to replace the conformation of fragment 81-122 using Rosetta as previously described^12, 13, 38^. In short, the dihedral angles from the donor segment are first recorded. The acceptor segment is mutated to poly Alanine and the number of residues are altered to match the length of the donor segment. A main-chain cut is randomly introduced in the acceptor segment and the dihedral angles previously recorded from the donor segment are applied. Following this step, a Cyclic Coordinate Descent (CCD) algorithm is applied to bring the two parts of the segment together and optimize the donor conformation. Following the conformation replacement, we proceed to the sequence optimization phase, utilizing RosettaDesign. This method relies on the Rosetta energy function and Monte Carlo sampling to generate sequences optimized for stability with the newly derived conformations.

### Design Strategy 2

The second approach, simpler in its implementation, involves creating sequence chimeras by replacing the target segment’s amino-acid sequence with the donor segment sequence and modeling the resulting structure using AlphaFold2 (see methods). Subsequently, we again used RosettaDesign to optimize the sequence.

### Design Strategy 3

Follows the same methodology as Strategy 2, but with increased complexity, by incorporating an additional conformation change within the segment spanning residues 159-194. This approach necessitates accommodating two independent conformational changes within the same protein scaffold, and managing any potential interactions between both segments.

### Design Strategy 4

Akin to Design Strategy 3, introduces two concurrent segment changes. However, it diverges in the execution of the design strategies. Firstly, we employ AlphaFold to model the new conformations. Secondly, the recently developed ProteinMPNN model is utilized for sequence optimization of the newly generated conformations.

### Soluble Expression Classifier

Given that both ProteinMPNN and RosettaDesign have been primarily refined for sequence recovery, they do not intrinsically optimize for high-yield heterologous expression. As such, the final scores assigned by each method to the resulting designs do not necessarily correspond to the probability of robust expression. Given this, we sought to develop an independent classifier trained specifically on the task of predicting soluble protein expression in *E.coli*.

Several machine learning models have been developed to predict protein solubility in various organisms, demonstrating varying degrees of accuracy. One particularly notable model, GraphSol, achieved state-of-the-art performance on the eSol dataset — A solubility database of ensemble *E.coli* proteins ^27^ . However, GraphSol is computationally intensive, requires additional evolutionary information (generated on the fly) and does not take into account high resolution structural information.

Our solubility classifier, the architecture of which is detailed in the Methods section, circumvents these limitations. In brief, the classifier constructs a graph representation of the Alphafold protein structure, wherein the amino acids are treated as vertices and an edge is defined between two vertices if the distance between them is less than 5Å. This classifier is an ensemble of 5 models wherein each of the models is a graph neural network composed of two graph attention layers activated by a Rectified Linear Unit (ReLU) function. Following attention, the graph representation is pooled via a Graph Multiset Transformer pooling operation. A final linear layer, followed by a sigmoid activation function, completes the process.

Our classifier exhibits exceptional precision and accuracy on a hold-out dataset, not utilized during training and validation, aligning with the performance of the current state-of-the-art classifier, GraphSol (Fig. 3; AUC=0.86 and Pearson’s R² = 0.41 when compared to the normalized expression values documented in the eSol dataset). Notably, our classifier demonstrates high computational efficiency, with the entire pipeline requiring between 10-30 seconds per protein sequence (depending on length) on a p2.xlarge AWS instance. This runtime includes the structure prediction step using the ColabFold implementation of AlphaFold.

**Figure 3.**
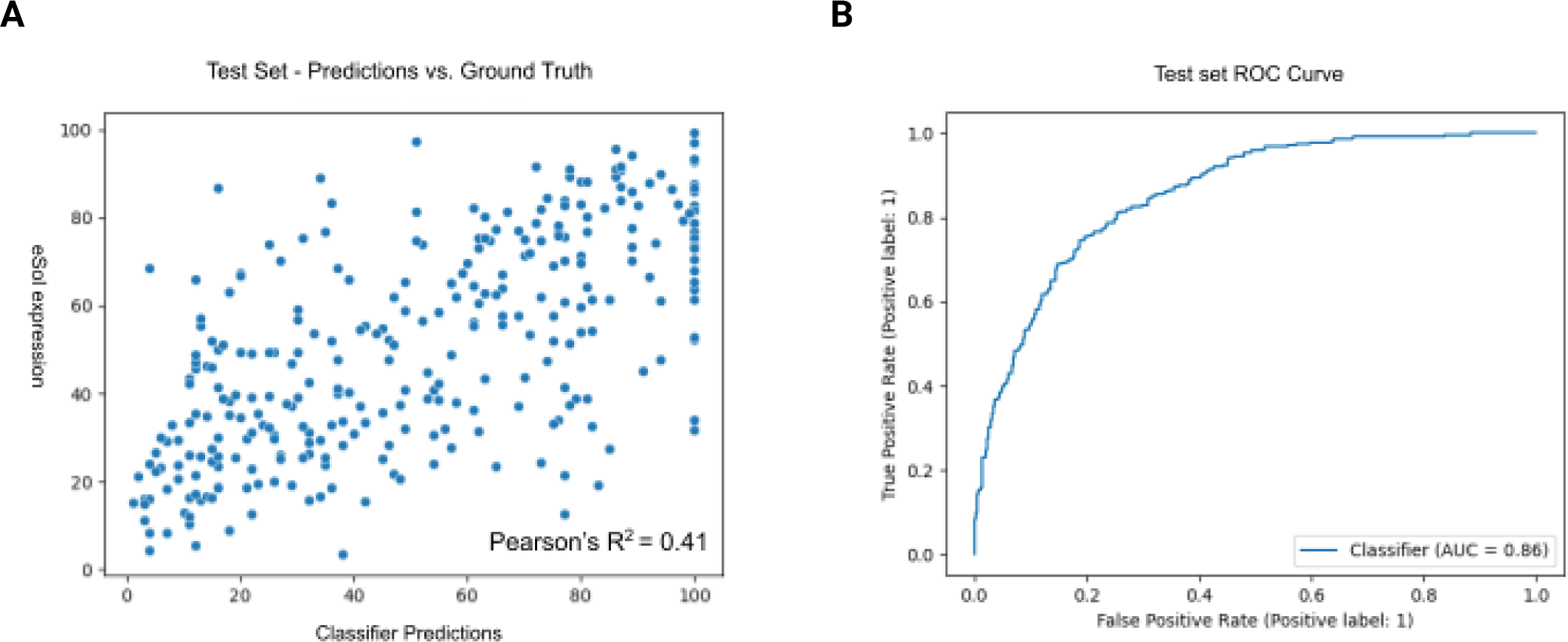
SolvIT Performance Metrics. (A) - Scatter plot showing the normalized soluble expression values of test-set entries in the eSol dataset as a function of classifier predictions scaled from [0,1] to [0,100]. (B) ROC Curve of classifier predictions on the test set vs. ground truth showing AUC of 0.86

The full CoSaNN protocol is depicted in Figure 4.

**Figure 4.**
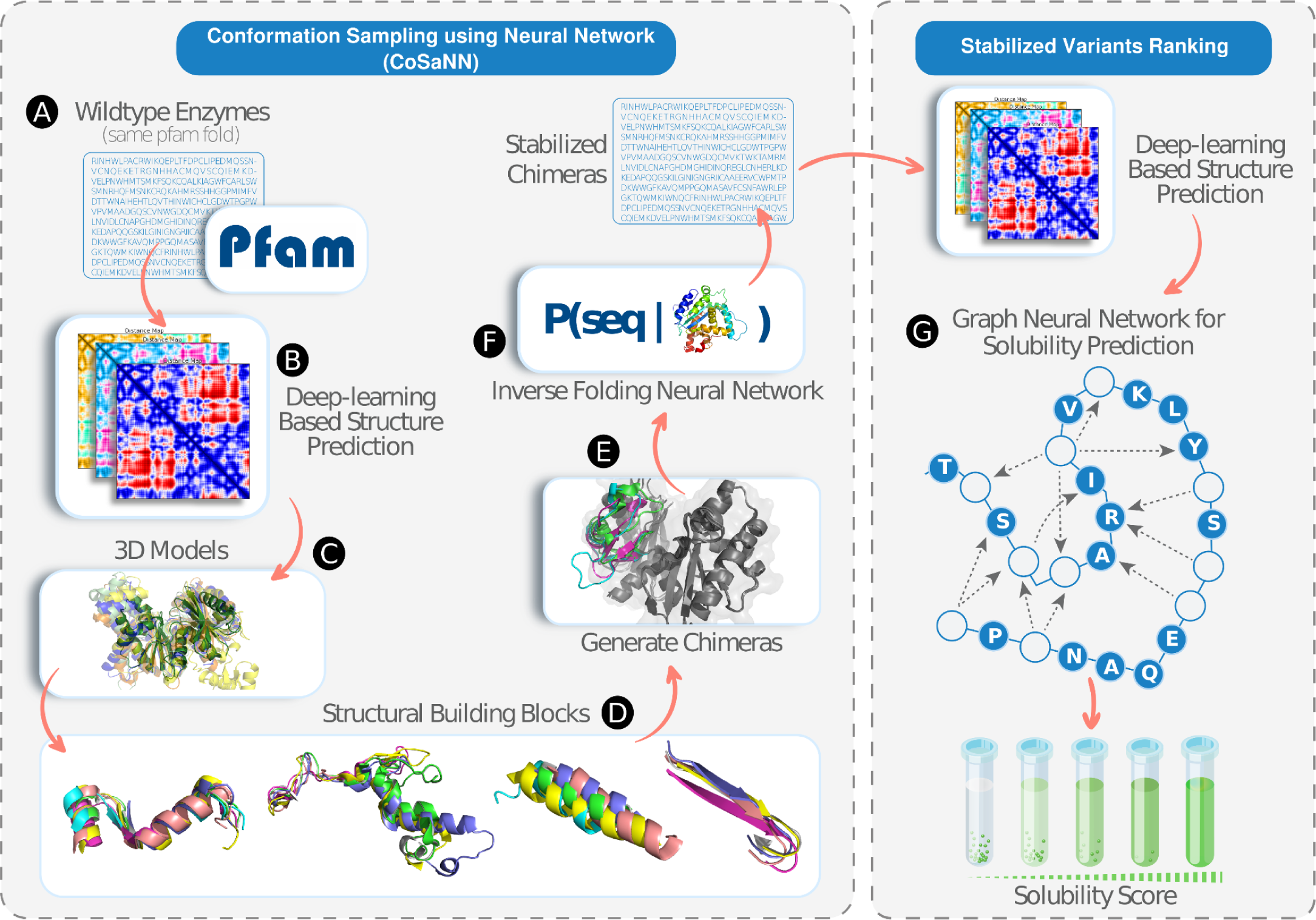
The CoSaNN Design Pipeline. (A) Protein family sequences are sourced from PFam and subjected to clustering, ensuring a 50% sequence identity. (B) Alphafold2 is employed to model the structure of each protein. (C) The resulting 3D models of protein structures are aligned. (D) Desired insertion points of each protein are identified, and relevant fragments are extracted from the aligned stem residues. (E) The extracted sequences replace the respective wildtype segment, resulting in the creation of chimeric proteins. These chimeras are subsequently modeled using Alphafold2 once more. (F) Inverse folding neural networks (or Rosetta) are utilized to optimize the sequence of each chimera. (G) Finally, SolvIT, a neural network designed to predict protein expression, is used to score each stabilized chimera.

#### Sequence analysis and structural diversity of the generated designs

Structural analysis of the design models demonstrates substantial structural and sequence diversity. The grafted segment encompassing residues 81-122 displays a Root Mean Square Deviation (RMSD) range from 0.45Å to 8.7Å, with a mean of 3.9Å in relation to the template segment (calculated over the backbone atoms). Similarly, the second grafted segment, spanning residues 159-194, exhibits an RMSD range between 0.62Å to 6Å, averaging at 2.66Å .

In terms of sequence diversity, the designs vary between 56% and 89% compared to the template enzyme. This range translates into more than 100 mutations relative to the template enzyme, indicating the breadth of sequence space sampled by our method.

Figure 5 summarizes the distribution of sequence identity and RMSD, sorted by the strategy employed. Not surprisingly designs that were modified with multiple segments (Design strategies 3 & 4) demonstrated higher sequence diversity relative to the template enzyme, with an average sequence identity of 65% and 60% respectively. Design strategy 4, which uses ProteinMPNN for sequence optimization, demonstrated even higher sequence divergence relative to the template enzyme compared to designs optimized using RosettaDesign. We expected these results given that ProteinMPNN samples a larger sequence space^20^.

**Figure 5.**
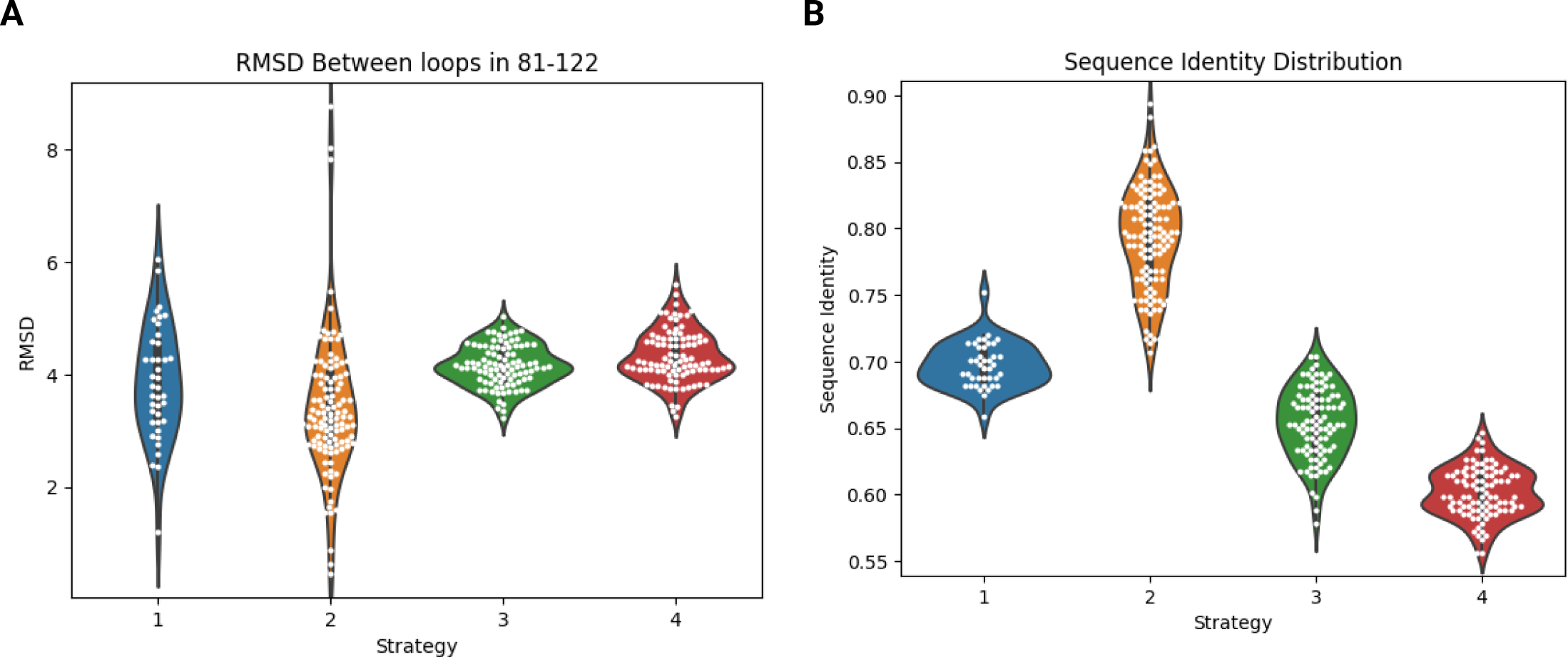
Structural and sequence diversity of generated designs grouped by strategy. (**A**) The distribution of RMSD between grafted segments and the original template conformation. (**B**) Distribution of sequence identity (over the entire protein sequence) for each of the designs relative to the template enzyme.

#### Soluble Expression and Thermal Melting Temperature Quantification

We cloned and expressed the designs along with two wild-type ROK family enzymes (ppgmk - the template used for all designs, and gmuE - an ATP-dependent fructokinase ^39^, serving as controls. Out of the 348 designs, 92 (26%) yielded more than 10µg total protein per 0.5ml induced culture volume, an additional 97 (28%) designs yielded detectable soluble protein amounts, and the remaining 165 (47%) designs showed no soluble expression. Analyzing the results in the context of the different design strategies utilized, we found that none of the designs produced using Design Strategy 1 (which involved backbone conformation sampling with Rosetta) showed detectable expression. In contrast, we observed soluble expression in 66% (78 out of 119), 33% (31 out of 95), and 85% (80 out of 94) of the designs created using Design Strategies 2, 3, and 4, respectively (summarized in Table 2).

**Table 2:** Summary of Enzymes’ Measurements Across Tested Strategies.

|  | Strategy 1 | Strategy 2 | Strategy 3 | Strategy 4 | Total |
| --- | --- | --- | --- | --- | --- |
| # of designs generated | 40 | 119 | 95 | 94 | 348 |
| Detectable Expression | 0 | 78 | 31 | 80 | 189 |
| Well Expressed (10µg total protein per 0.5ml induced culture volume) | 0 | 50 | 3 | 39 | 92 |
| T<sub>m</sub> > 56.7°C | 0 | 35 | 8 | 70 | 113 |
| Active (ATP+Glc) | 0 | 42 | 0 | 4 | 46 |
| Active (HexaP+Glc) | 0 | 54 | 2 | 0 | 56 |
| Active (ATP and HexaP) | 0 | 42 | 0 | 0 | 42 |

Interestingly, we noted a distinct difference in the expression rates between strategies 3 (RosettaDesign) and 4 (ProteinMPNN). Both strategies involve the simultaneous grafting of two different segments, with one of the grafted segments including of the Zinc finger motif which is absent from the ppgmk template ^40^. This outcome suggests that the simultaneous grafting of multiple segments, particularly when these separate sequence segments interact in the three-dimensional protein structure, necessitates a nuanced understanding of both the grafted segments and the accepting template within their specific structural context. These high-order interactions learned during the training of ProteinMPNN can be generalized to novel protein conformations, as is the case in this study, and aligns with previous findings^20^.

### Thermal Melting Temperature Quantification

To evaluate the thermal stability of the designs, thermal melting temperature (Tm) was measured using Differential Scanning Fluorimetry (DSF) with SyproOrange as a probe ^41^. 38% of all designs (a total of 135) underwent successful thermal stability evaluation, including the control wildtype enzymes ppgmk (the template enzyme) and gmuE, exhibiting a Tm of 56.7°C and 62.8°C, respectively.

Remarkably, 83% of the assessed designs (113 out of 135) demonstrated a higher Tm than that of ppgmk, the original template enzyme. Notably, Design Strategy 4, while being the most divergent in sequence and conformation relative to the template, yielded the highest average Tm among all tested proteins, with 70 (78%) designs surpassing the Tm of ppgmk (Fig. 6B).

**Figure 6.**
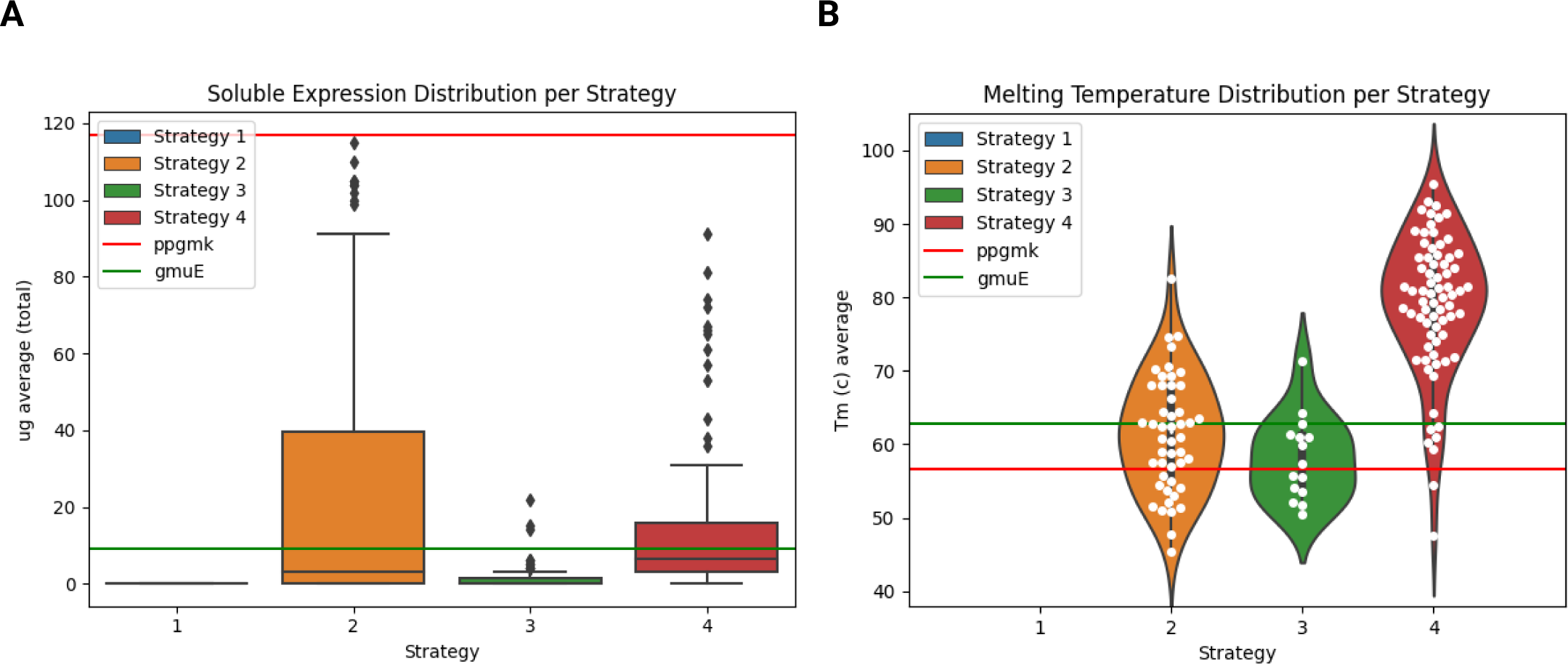
Assessment of Expression Rates, Thermal Stability Across Different Protein Design Strategies. (A) Comparison of the expression rates of different strategies. (B) Comparison of the thermal melting temperature of different strategies (Strategy 1 not shown due to lack of expressed variants).

These results are visualized in Figure 6, which presents a comprehensive distribution of the designs’ melting temperatures, grouped by design strategy and contrasted against the ppgmk and gmuE controls.

#### Solubility Classifier Evaluation on Backbone Designed Enzymes

Major backbone perturbations and large sequence deviations from the wild-type have the potential to destabilize the protein ^42^, increase its tendency to misfold, and hence, decrease its total soluble expression. Therefore, we wanted to test whether classification to soluble/insoluble expression can be explained by trivial parameters such as sequence identity or RMSD to the original template enzyme.

Each measured variable (sequence identity and RMSD for both modified segments) exhibited a modest, yet statistically significant, correlation with soluble protein yield. Specifically, sequence identity demonstrated a Pearson’s R^2^ value of 0.22 (p=2e-5), the RMSD value for the segment between residues 81-122 had an R^2^ value of -0.22 (p=4e-5), and the RMSD value for the segment between residues 159-194 exhibited an R^2^ value of -0.31 (p=1e-5). Those measurements however show poor binary predictability for whether the expression yield is higher than 10µg total protein per 0.5ml induced culture volume, achieving AUCs of 0.54,0.58,0.53 for sequence identity, segment 81-122 RMSD and segment 159-194 RMSD (Figure S1), respectively. These findings indicate that the likelihood of expression in yields higher than 10µg total protein per 0.5ml induced culture volume cannot be explained by simple observations like the mutation load or the degree of structural perturbations to the template structure, which motivated us to develop a new classifier which can capture more intricate interaction between protein conformation, sequence, and predicted expressibility. We evaluated the predictive capacity of our soluble expression classifier by forecasting the solubility of each respective design. Samples that yielded in excess of 10µg total protein per 0.5ml induced culture volume were denoted as positive, while those with lesser yields were categorized as negative. An ensuing ROC plot was drafted, and the area under the curve (AUC) was calculated, attaining a value of 0.79. Establishing a probability threshold greater than 0.7 yielded an accuracy of 70%, a Positive Predictive Value (PPV) of 43%, and a Negative Predictive Value (NPV) of 93%.

Earlier research ^43^ has suggested that both ESMv2 and ProteinMPNN, despite not being explicitly trained to predict protein fitness, thermal stability (Tm), or expression yield, are capable of offering reliable unsupervised predictions regarding these experimental properties. However, when these models were applied to our designs, their predictive capacity was mediocre. The ESMv2 mean likelihood on the design sequences achieved an AUC of 0.69, whereas the ProteinMPNN mean negative log-likelihood scores realized a classification performance with an AUC of 0.66.

When using our classifier to predict the solubility of designed enzymes that were not part of the classifier’s training dataset, the calculated PPV suggests that experimentally testing designs with a prediction score higher than 0.7 would result in over 50% of the designs being well expressed, compared to 26% when selecting designs generated using the same design pipeline but without the additional solubility classification step.

#### Enzyme Activity Measurement

Designing enzymatic activity requires sub-angstrom precision in the configuration of catalytic residues with respect to each other and the substrate. Therefore, the ability to create active enzymes serve as strong evidence for the accuracy of our design method.

We screened the soluble designs for glucokinase kinase activity. We incubated the enzymes with glucose and either ATP or hexametaphosphate for 16 hours and measured the concentration of glucose-6-phosphate created using a modified picoProbe assay as described in the Methods section. In total, 60 designs were active with either hexametaphosphate or ATP; 54 of the active designs were created using Design Strategy 2 (90% of active enzymes), 2 designs were created using Design Strategy 3 (3.3% of active enzymes), and 4 designs were created using Design Strategy 4 (6.6% of active enzymes). Interestingly, 8 (13%) designs showed obligatory polyphosphate dependent activity while the template enzyme can utilize both ATP as well as inorganic polyphosphate. Obligatory polyphosphate activity is not common in natural enzymes and has been documented in a handful of cases ^40, 44^. The emergence of this feature in our designs underscores the potential of our method to introduce novel, non-natural enzymatic traits.

We selected 2 designs with high soluble expression rates, Tm and activity (design ids: SKFe, 1rvg, both from Strategy 2) for additional biochemical and biophysical characterization. We decided to use 1rvg as our reference for activity measurements. We quantified its maximal activity as 100%, corresponding to the amount of substrate 1rvg synthesized at a temperature of 25°C within a span of six minutes. We then measured the relative activity of each of the designs and compared it to the ppgmk template, as a function of reaction temperature (Fig. 7A). Both designs show overall higher activity than the ppgmk template across all measured temperatures above 35°C, with SKFe retaining close to 60% relative activity at 85°C and more than 40% relative activity at 95°C for the measured time duration.Conversly, The ppgmk template enzyme loses activity completely at 95°C.

**Figure 7.**
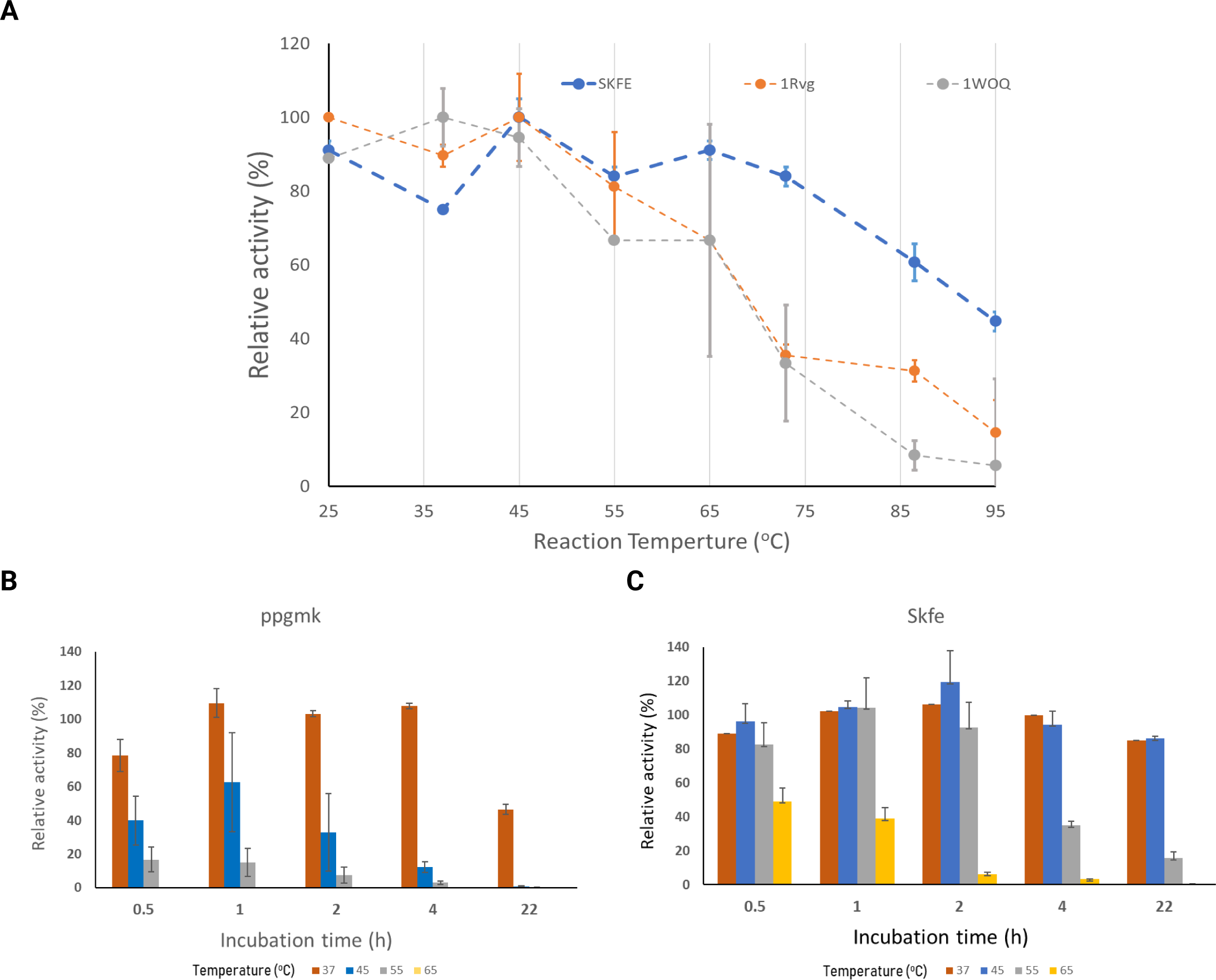
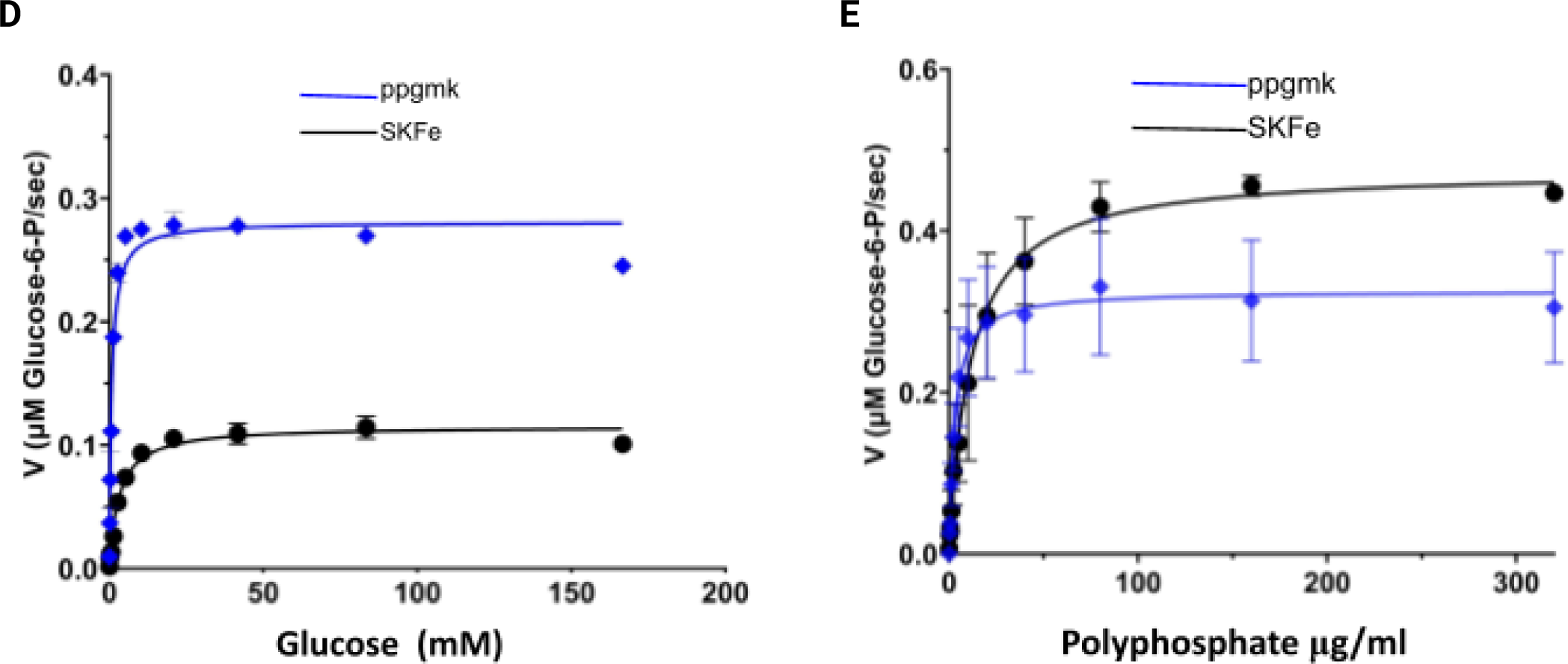
Enzymatic activities of selected enzymatic design. Each measurement was performed in duplicates. (A) - The glucose phosphorylation reaction in different temperatures for SKFe, ppgmk and 1rvg. (B) + (C) - Thermal stability - of ppgmk and SKFe activity as a function of incubation time and temperature. (D) - SKFe and ppgmk Michaelis-Menten kinetics on glucose. (E) SKFe and ppgmk Michaelis Menten kinetics on hexametaphosphate. Error bars represent the standard deviation of two independent experiments. The datasets were significantly different according to the extra sum of squares F-Test (P=0.006, n=2).

We next compared the capability of SKFe and the template enzyme, ppgmk, to sustain catalytic activity after prolonged temperature stress in different temperatures, by measuring the relative remaining activity after several incubation times and temperatures (Fig. 7B, See Methods for additional details).

Briefly, the enzymes were incubated without their respective substrates at varying temperatures for distinct time intervals, after which they were cooled to 37°C. Subsequently, substrates were introduced and product formation was measured after a duration of six minutes. In contrast to ppgmk, which became inactive after 4 hours at 55°C and after only 30 minutes at 65°C, SKFe retained over 80% of its activity even after 22 hours at 55°C, and more than 40% after 30 minutes at 65°C.

We proceeded to characterize and compare the kinetics of SKFe and ppgmk using polyphosphate and glucose as substrates at 37°C (Fig. 7D,7E). We calculated the Km of SKFe to be 3.15mM with respect to glucose and 2.9μg/ml with respect to polyphosphate. The equivalent parameters for ppgmk were calculated to be 0.75mM and 12.26μg/ml respectively. We hypothesized that SKFe’s higher affinity towards polyphosphate may come at the cost of lower affinity to ATP as SKFe had no detectable activity when ATP was used as the phosphate donor.

## Discussion

Enzyme design presents one of the last remaining frontiers in computational protein design, necessitating the optimization of multiple objectives, from expression, stability, conformational plasticity and catalytic function. The last being particularly challenging as it requires the modeling of quantum phenomena that for the most part still remains poorly understood. The ability to design new-to-nature enzymes not only poses a stringent test of our understanding of the ‘protein rule set’ but also presents a transformative technology as it will enable new manufacturing processes, orders of magnitude more efficient than current ones.

While designing completely *de novo* catalytic machinery still presents a daunting challenge in computational protein design, here we present an approach that circumvents this challenge and focuses on expanding the catalytic repertoire of natural enzymes by taking inspiration from evolutionary processes which generate diversity by conserving the core catalytic machinery while introducing changes to peripheral conformations and sequences. While this engineering concept by itself is not new and is routinely used in experimental techniques such as directed evolution, these techniques are extremely inefficient in exploring the inconceivably large structure-sequence space. Natural evolution explores the structure-sequence space in a stepwise manner. This conservative approach reflects a biological constraint: introducing too many mutations at once to a protein increases the risk of generating a non-foldable or non-functional protein.

The advent of computational design tools like Rosetta have enabled the ability to ‘leap frog’ through the structure-sequence space allowing for a more expansive exploration. However, these techniques also have their shortcomings. The reliance on random moves to explore the protein space and the use of a limited scoring function that fails to capture high-order amino-acid cooperativity, means that many design trajectories that explore the protein space end up in non-foldable regions.

In this work we demonstrate a novel approach that leverages the latest advances in neural networks and directly addresses these challenges. We treat the structure-sequence design process as two separate steps. First, we exploit the ‘forced-to-predict’ feature of trained structure prediction deep neural networks, meaning that even for input sequences that are ‘unfoldable’ in the real world, these models would still produce a ‘native like’ protein conformation. Following this step we are most likely in a region of sequence-structure space that is non-foldable (Fig. 8). To rectify this, we apply an inverse folding neural network to move us along the sequence dimension while keeping the conformation fixed (Fig. 8), putting us back in a foldable region of the protein space. Lastly, we apply an additional optimization layer to increase the probability of generating highly expressible designs. This process simultaneously dramatically increases protein space samping efficiency while at the same time greatly simplifying it.

**Figure 8.**
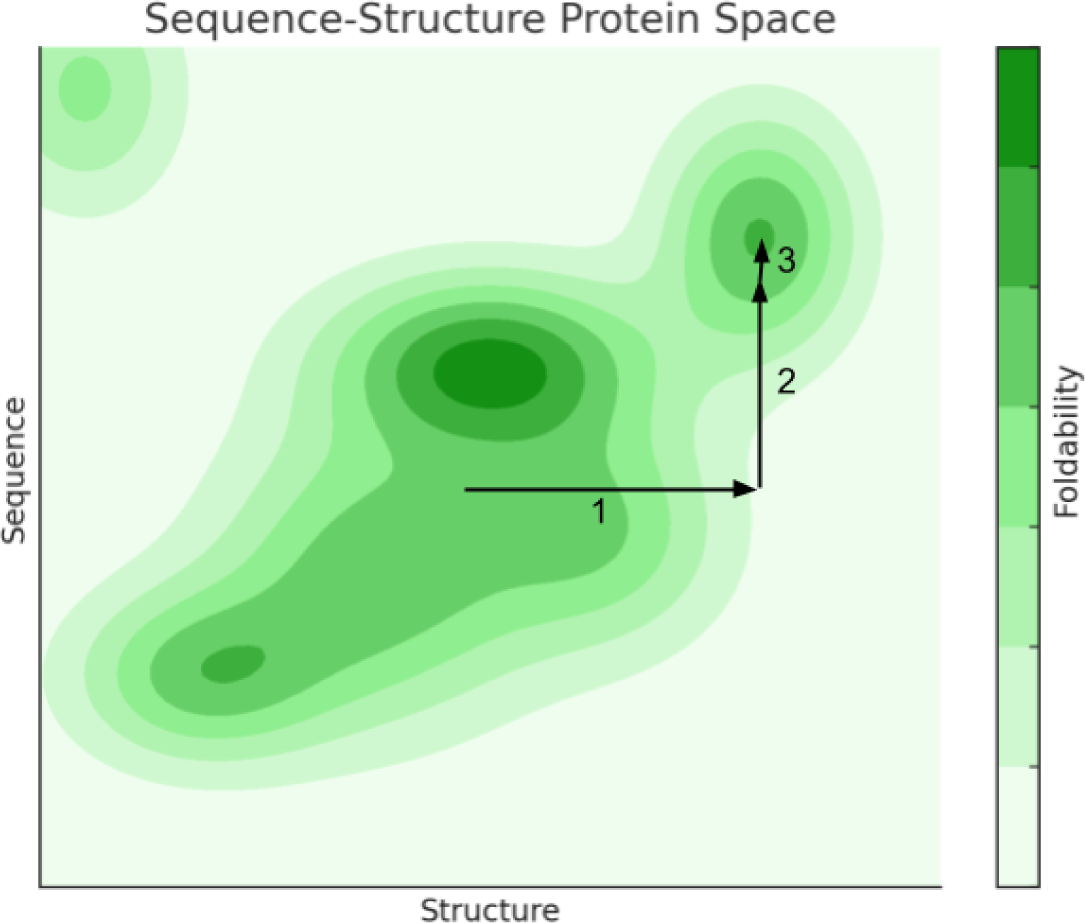
CoSaNN traversal of the sequence-structure space. The contour map represents a 2D projection of the possible sequence-structure protein space. Darker regions represent regions with high foldability. During the CoSaNN design flow (**1**) we first move along the Structure coordinate using protein structure prediction NN. This step potentially places us outside a foldable region.(**2**) The next step uses an inverse folding NN to move back to a foldable region along the same Structure coordinate.(**3**) Lastly, we apply SolvIT to optimize for expression within the foldable region.

Using this approach, CoSaNN is able to generate novel enzymes that, while being extremely divergent from natural homologues in conformation and sequence space, still maintain high activity and demonstrate superior stability and expressibility, which are fundamental properties for any commercially relevant enzyme. Not only that, but we directly tackle the challenge of designing enzymes with elaborate activity mechanisms that undergo significant conformational changes upon substrate binding, underscoring our ability to capture intricate amino-acid interactions. This ability has implications for other protein design challenges such as allosteric regulation.

Our ability to dramatically modify an enzyme’s structure without adversely affecting neither the catalytic activity nor the allosteric mechanism enables us to encode new substrate specificity to a much larger extent than what is possible through single point mutations.

Lastly, we have shown that by adding an expression filter layer to our method we can generate designs with scores of mutations, far beyond what can be achieved using currently available experimental techniques, we did not have to rely on high throughput techniques to screen our enzyme designs which dramatically reduces the complexity and time required to develop new enzymes.

## Supplementary Material

**Figure S1:**
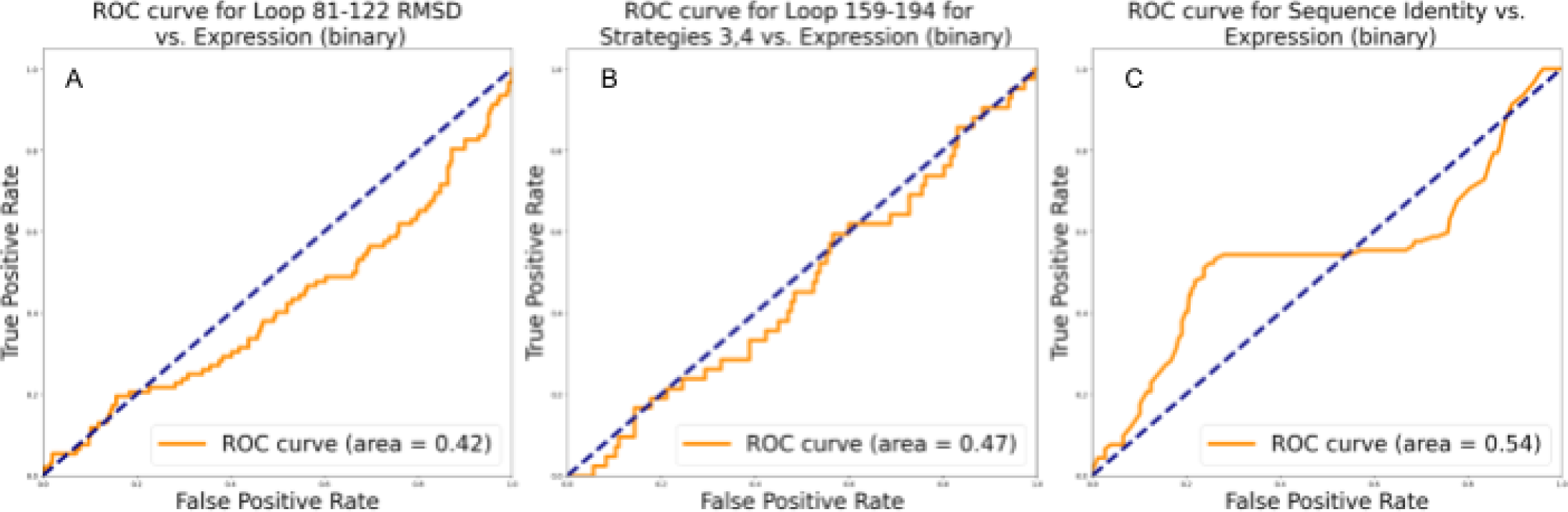
ROC Curves Illustrating the Predictive Performance of Sequence and Structural Diversity Features for Expression Rate. **(A)** RMSD of loop 81-122 to the wildtype loop of ppgmk as a predictor of expression rate. **(B)** RMSD of loop 159-194 (for strategies 3,4) to the wildtype loop of ppgmk as a predictor of expression rate **(C)** Sequence identity of designs to ppgmk as a predictor of expression rate

Table S1: Protein Expression, Tm and activity data (Separate File)

